# On the molecular mechanism of SARS-CoV-2 retention in the upper respiratory tract

**DOI:** 10.1101/2020.07.29.227389

**Authors:** Kristina A. Paris, Ulises Santiago, Carlos J. Camacho

**Affiliations:** Department of Computational and Systems Biology, University of Pittsburgh, Pittsburgh, PA, 15260, USA

## Abstract

Cell surface receptor engagement is a critical aspect of viral infection. At low pH, binding of SARS-CoV and its ACE2 receptor has a tight interaction that catalyzes the fusion of the spike and endosomal membranes followed by genome release. Largely overlooked has been the role of neutral pH in the respiratory tract, where we find that SARS-CoV stabilizes a transition state that enhances the off-rate from its receptor. An alternative pH-switch is found in CoV-2-like coronaviruses of tropical pangolins, but with a reversed phenotype where the tight interaction with ACE2 is at neutral pH. We show that a single point mutation in pangolin-CoV, unique to CoV-2, that deletes the last His residue in their receptor binding domain perpetuates this tight interaction independent of pH. This tight bond, not present in previous respiratory syndromes, implies that CoV-2 stays bound to the highly expressed ACE2 receptors in the nasal cavity about 100 times longer than CoV. This finding supports the unfamiliar pathology of CoV-2, observed virus retention in upper respiratory tract^1^, longer incubation times and extended periods of shedding. Implications to combat pandemics that, like SARS-CoV-2, export evolutionarily successful strains via higher transmission rates due to retention in nasal epithelium and their evolutionary origin are discussed.

---

Although accurate assessments are still evolving, reports from the world health organization indicate that infection with SARS-CoV-2 (CoV2) is significantly different from infection with previous respiratory viruses. The biggest distinctions are a longer incubation period and increased viral shedding (both likely responsible for higher rates of transmission), strong correlation of infected-fatality rates (IFR) with age and comorbidities, higher IFR for males relative to females, and minimal impact in young children. While this complex pathology could be due to complex genomic changes triggered by the virus in cells, tissues, and organs^2^, longer incubation and infectivity time scales also suggest that differences could have a biophysical origin.

Work on SARS-CoV (CoV1) has already determined that the virus enters cells via receptor-mediated endocytosis in a pH-dependent manner^3^ that is characterized by co-translocation of the viral spike glycoprotein and its specific functional receptor, the angiotensin-converting enzyme 2 (ACE2), from the cell surface to early endosomes. Key steps that control the fate of the virus in the early and late endosome are driven in part by lowering the pH from 6.5-to-6.0 and from 6.0- to-5.0^4^, respectively; exposure to low pH triggers a spike catalyzed fusion between the viral and endosomal membranes followed by viral genome release. The cell machinery is then hijacked to replicate and assemble new virus particles that eventually exit the cell, primarily through budding. Broadly speaking, this is the same pH-dependent endocytic path followed by the influenza virus^5,6^, and likely all other coronaviruses (including MERS-CoV).

Cell surface receptor engagement is a critical aspect of viral infection and life cycle. While infections by CoV1 and CoV2 are mediated by the ACE2 receptor, MERS-CoV (MERS) gains entry to cells through dipeptidyl peptidase (DPP4)^7^. Fig. 1 shows structures for the receptor-binding domains (RBD) in complex with their receptors for all three viruses. Surprisingly and likely significantly, while the RBD of CoV1^8^ and MERS^9^ have one Histidine on opposite ends of their binding interface, CoV2^10^ does not have His residues in this domain. As we have been able to determine to date, all known strains of CoV2 have mutated away their last His residue that is still present in the RBD of CoV1/MERS and related zoonotic viruses (see Table 1). His protonation is important in the low endosomal pH for spike fusion with the membrane, however, little is known on the purpose of His in neutral pH that is relevant for the virus to establish an initial foothold in the respiratory system. Here we set to decipher the mechanistic role of the remaining His residue that distinguishes the RBDs of CoV1 and CoV2. We discovered that deletion of the pH-switch in the RBD of CoV2 results in a tighter bond to ACE2 than CoV1, retaining the virus in the nasal cavities for significantly longer time scales. More interestingly, we discover that this mechanism is fully reconciled by a single point mutation in the sequence of related coronaviruses in Sunda pangolins^11^.

**Figure 1.**
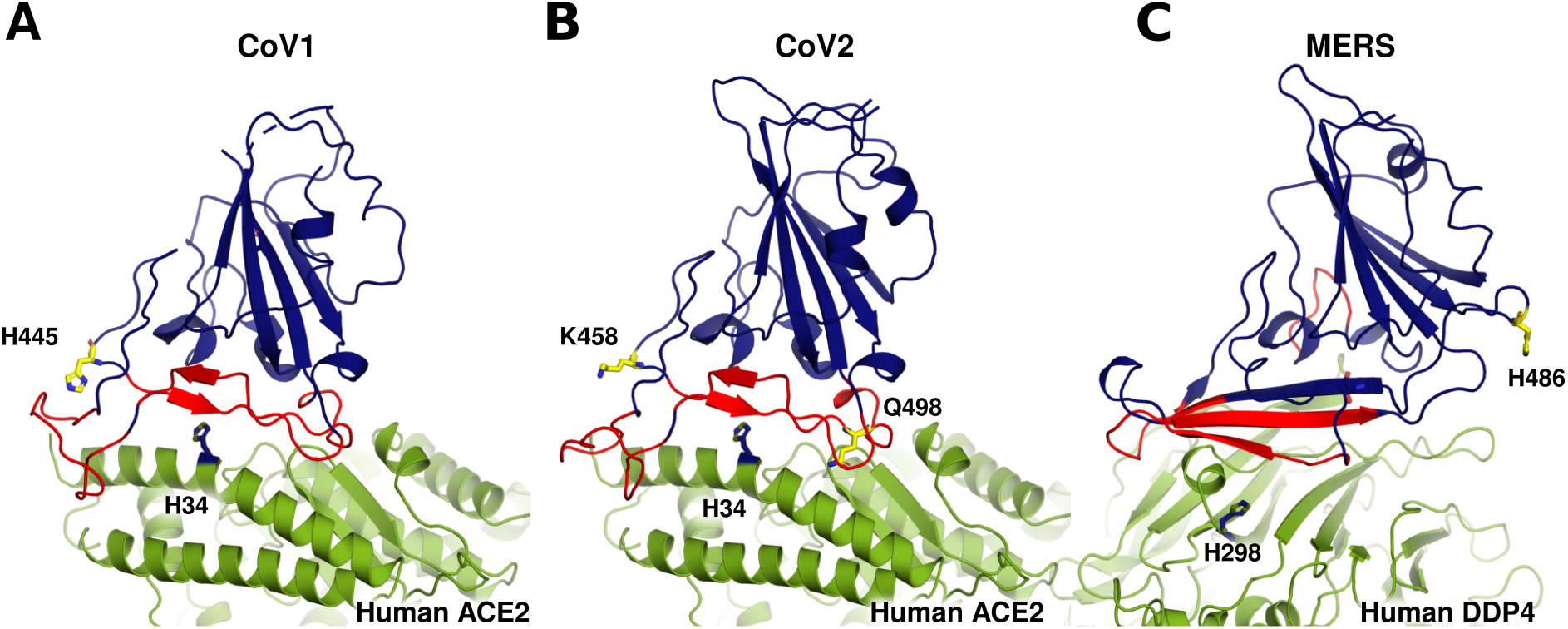
Deletion of Histidine residues in receptor binding domain (RBD) of CoV2. Co-crystal structures of the RBD (blue) and their specific receptor (green) for (A) SARS-Cov (PDB 2AJF; CoV1), (B) SARS-CoV2 (PDB 6LZG; CoV2) and (C) MERS-CoV (PDB 4L72; MERS). RBD binding interface is shown in red. RBD’s of Cov1 and MERS have one histidine each (yellow), H445 and H486, respectively. Key residues K458 and Q498 in CoV2 are indicated in yellow. H34 (in blue) is the only histidine in the binding interface of ACE2.

**Table 1.**
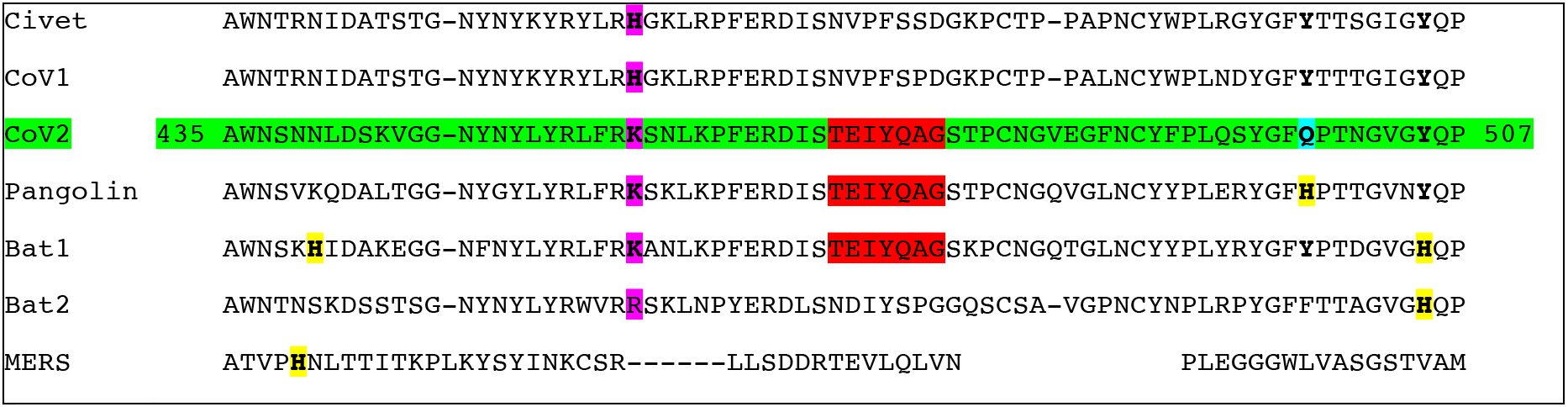
Sequence alignments of pH domains in RBD CoV2 and related viruses. Histidine (H) mutation in CoV1 to K in CoV2 is highlighted in pink column. Conserved motif relevant for interaction with K458 are shown in red for CoV2, Pangolin, and Bat1. H residues that co-localize with MERS pH-switch are shown in yellow. Only Histidine in RBD of pangolin-CoV corresponds to Gln (cyan) in CoV2. NCBI Locus AAU04646 (Civet); 6CRV_C (CoV1); 6VSB_A (CoV2); QIQ54048 (Pangolin); QHR63300 (Bat1); AGZ48806 (Bat2); QBM11748 (MERS).

## pH-switch reveals distinct binding landscapes

We studied the role of pH in the RBD of the spike proteins by performing three independent 500 ns unconstrained molecular dynamics simulations (MDs) of CoV1 (PDB 2DD8^12^) and CoV2 (PDB 6LZG^10^) at both physiological and low pH ~ 5.0 conditions (See Methods). At neutral pH, the equilibrium binding affinity of the RBD of CoV2 and ACE2 has been shown to be slightly stronger than that of CoV1^13,14^. No such measurements exist at low pH, which in practice should protonate His residues from neutral to positively charged (His^+^) by the addition of an extra hydrogen. Of note, although not studied here, the RBD of MERS-CoV has previously been shown to have two distinct conformations at neutral and low pH^15^.

At neutral pH, MDs revealed important differences between the RBD of CoV1 and CoV2. For CoV1, the dynamics of the binding interface (Fig. 2A) is divided between (a) a loop (F_460_SPDGKPCTPPALNCY_475_) that displays not-bound-like conformations (~ 5.8 Å), and (b) the remainder of the binding interface that consistently adopts bound-like conformations (~ 1.1 Å) relative to its co-crystal (PDB 2AJF). Simulations for tautomer NεH at His445 are shown in Fig. S1. Remarkably, pH-independent CoV2, which mutates the last Histidine, His445, in CoV1 to Lys458 in CoV2, yields almost exclusively bound-like conformations for the whole binding interface (Fig. 2B).

**Figure 2.**
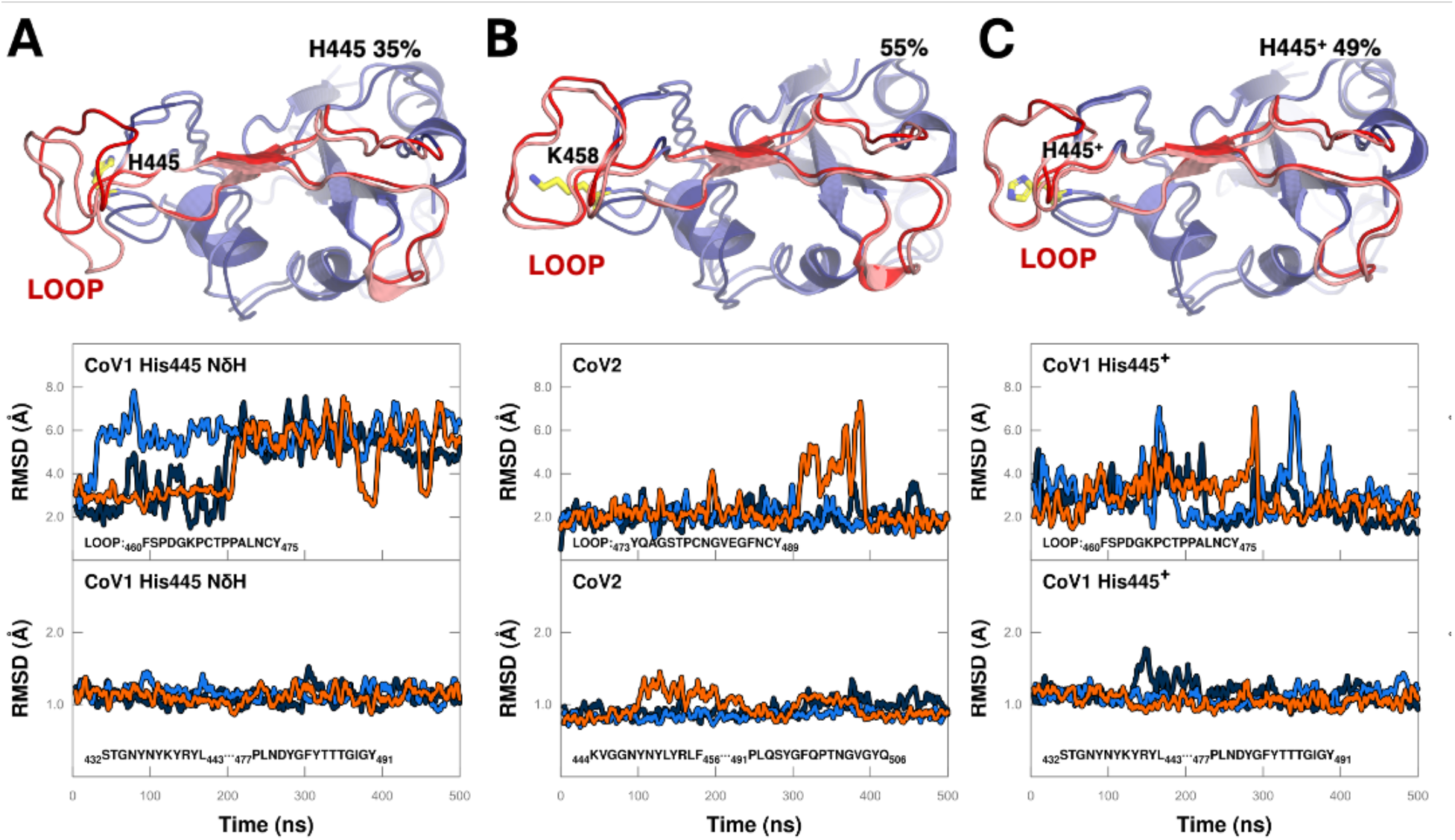
Role of pH switch in CoV1 relative to CoV2. Overlap of binding interface of RBD co-crystal (blue) and representative MD snapshot (red) corresponding to the centroid from largest 1.8 Å cluster in last 300 ns of the MD (% of cluster size relative to simulation time is indicated); Three independent MDs showing RMSD relative to co-crystal of (a) 16 amino acid loop (or its homolog) as a function of time, and (b) the remaining of the binding interface (or its homolog) that stays in a bound-like conformation for 100% of the simulation time, for (**A**) CoV1 at neutral pH (His445), (**B**) CoV2, and (**C**) CoV1 at low pH (His445^+^). Loop in CoV1 switches between not-bound-like (~ 5.8 Å) to bound-like (~ 1.9 Å) for His445 and His445^+^, respectively. Bound-like behavior of His445^+^ resembles CoV2.

Of note, independent molecular dynamics of neutral pH CoV1 and CoV2 RBD bound to ACE2 had been previously performed by the D. E. Shaw Research group.^16^ Consistent with our unbound MDs, the loop in CoV1 is found to move away from the co-crystal structure, sampling not-bound-like conformations significantly more than the homologous loop in CoV2. As in Fig. 2, the remainder of the binding interface (not including the loop) is stable around the bound-state throughout the 10 *μ*s MDs.

Low pH is critical for the activation of the spike fusion with the endosomal membrane^17^. Hence, one would expect that both CoV1 and CoV2 interaction with ACE2 to be similar. Strikingly, our MDs showed that a protonated His445^+^ in Cov1 (Fig. 2C) switches the dynamics of the loop from not-bound-(~ 5.8 Å) to bound-like (~ 1.9 Å), establishing the same phenotype as the one observed in CoV2. As shown in the representative Supplementary Movies 1, the interactions of His445^+^ relative to His445 with the switching loop are more stable, and resemble Lys458 that consistently engages seven conserved nearby residues T_470_EIYQAG_476_ (Table 1) to stabilize the bound-like phenotype. Low pH should also protonate His34+ in ACE2 (Fig. 1). However, His34^+^ should only increase the overall binding affinity for both CoV1 and CoV2 since their corresponding co-crystal structures suggest that they should form the same intermolecular hydrogen bond network (Fig. S2).

Implications of these findings in the binding free energy landscape of CoV1 and CoV2 are sketched in Fig. 3. Namely, at low pH, the similar dynamics and affinity of both CoV1 and CoV2 with their specific ACE2 receptor is drawn as a deep well in the binding landscape. This reflects the fact that breaking the complex should entail detaching all the contacts at once since the full binding interfaces for both viruses are stabilized around the bound conformation before and after binding ACE2. However, at neutral pH the binding landscape of CoV1 is different due to the pH-switch in its RBD. Specifically, Fig. 2A reveals a higher free energy transition state characterized by a bound-like behavior for most of the binding interface except for a distal loop that switches to a not-bound-like state. Of note, equilibrium thermodynamics alone cannot distinguish between the different landscapes shown in Fig. 3. In fact, the presence of a transition-state can only be revealed by studying the dynamics of the interaction.

**Figure 3.**
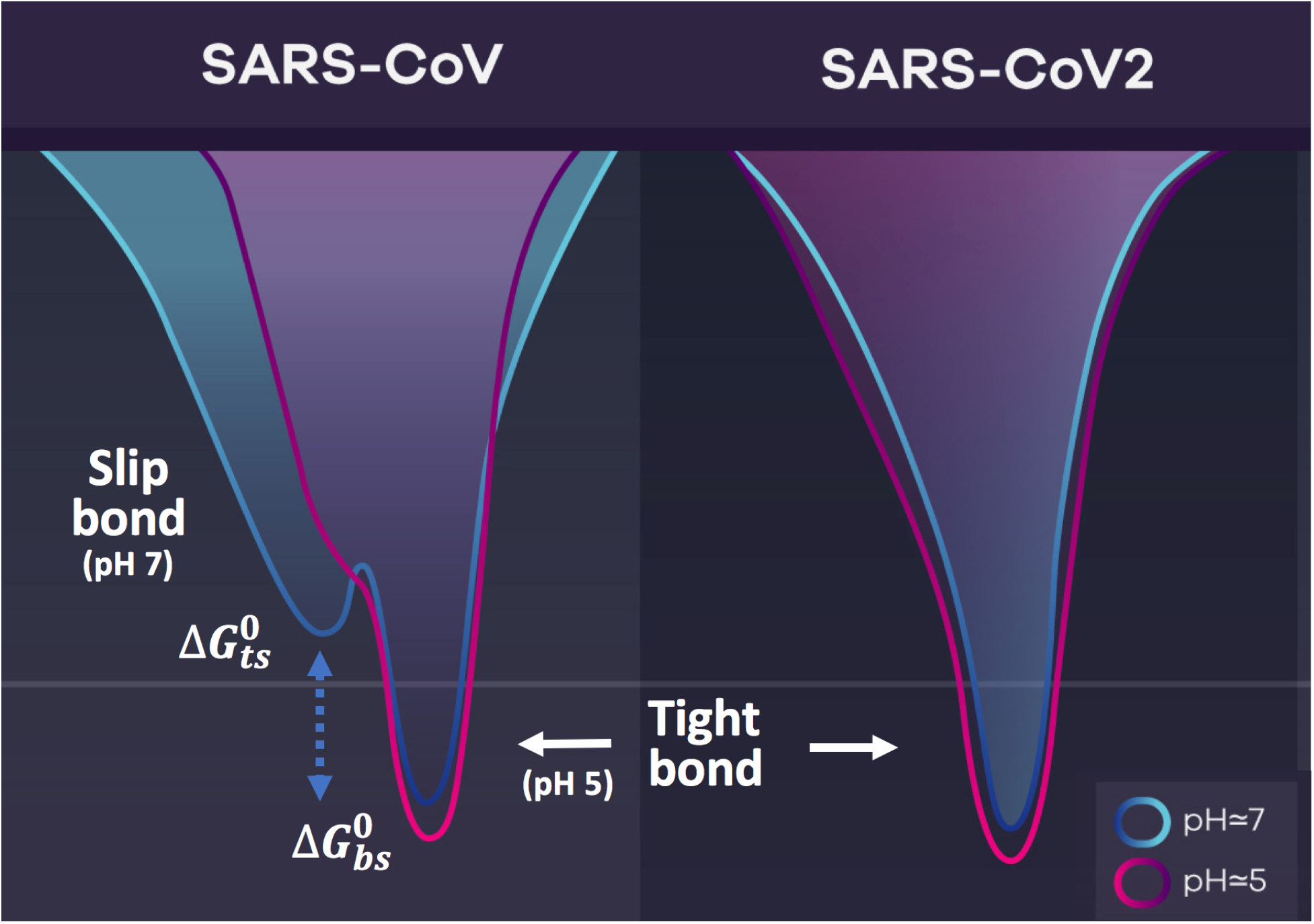
Free energy binding landscapes of RBD of SARS-CoV1 and SARS-CoV2 and ACE2 receptor. At neutral pH, CoV1 dynamics reveals a transition-state that gives rise to a slip-bond in the presence of external forces such as shear flow. At low pH, this state is eliminated, and similar to CoV2, the landscape is depicted as a deep well corresponding to a tight-bond. Low pH curves are shown with lower binding free energies to reflect the role of His34^+^ in ACE2.

## Slip-bond enhances CoV1 off-rate

What are the implications of this transition state found in CoV1 but not in CoV2 at neutral pH? To answer this question, we need to consider that viral particles in the respiratory system are under small external strains from, e.g., the ever-present shear flow in the respiratory airways, when engaging cell surface receptors. This shear flow generates tensile forces that pull and push on the chemical bonds between the RBD and ACE2 shifting their behavior from equilibrium to out-of-equilibrium kinetics. Specifically, while the association rate of CoV1 and ACE2 was found to be diffusion-limited with 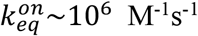, the off-rate cannot be assumed to correspond to the equilibrium formula 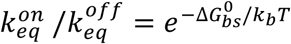 as stated in, say, Shang et al.^14^. Instead, one needs to independently measure the lifetime of bonds under tensile forces.

This problem is well understood both theoretically^18,19^, and experimentally^20,21^. Based on Arrhenius theory, the rate constant for detachment when a “spring-like” force is applied to the bond can be written as^18^

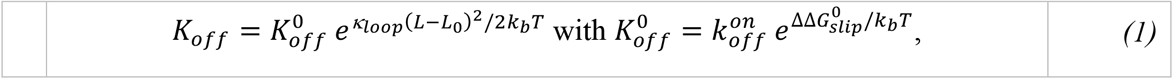

where 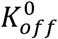 is the baseline rate of detachment; 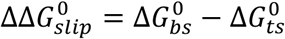, with 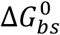 and 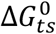 correspond to the binding free energy of the bound-state and transition-state (Fig. 3), respectively; and, *k*_*loop*_ is the “spring” constant associated with the tensile forces exerted on the loop domain at a given gap-width *L*. Note that in the absence of a transition-state, this rate corresponds to the classical 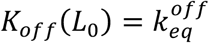 where *L*_0_ is the equilibrium (no tensile forces applied) intermolecular gap length.

It is clear that the spring approximation is exact for small deformations and in quasi-equilibrium conditions where the ramp rate for the force must be slow. In general, shear flow will twist around its long-axis to reach the transition state from the bonded state. And, since *k*_*loop*_ > 0, whether pulling or pushing on the chemical bond, this motion always loosens “tightness”. This corresponds to a typical slip-bond^19–21^, whose lifetime is shortened by tensile forces acting in the bond.

## Off-rates of CoV1/CoV2 detachment from ACE2

Based on the binding landscape revealed by our simulations (Fig. 3), the equilibrium binding free energies 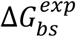 for these complexes^13^, and a partition of the binding free energy between (a) the loop and (b) the remainder of the binding interface computed using the server *FastContact*^22^, we can estimate the baseline rate of detachment for the RBD/ACE2 bond under tension 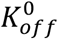 in Eq. (1) for (i) the slip-bond to the transition-state (ts) of CoV1, (ii) full detachment from this state, and (iii) full detachment from the tight-bond of CoV2 (Table 1). The key finding of this analysis on the dynamics of virus engagement with cell surface receptors is that, at neutral pH, CoV1 stabilizes a transition-state, which detaches a loop that is estimated to contribute about one fifth of the total binding free energy or about 2.6 *kcal/mol*. As a result, small external strains applied by the shear flow in the respiratory system trigger a “slip-bond” that enhances the off-rate *K*_*off*_ of the viral particle by a factor of 80 relative to 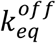; and, 100 times faster than the tight-bond of CoV2. Of note, alternative binding free energy measurements have suggested a 10-fold weaker binding14 than in Table 2. These differences will only re-scale 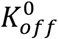 by a factor of 10 or so but will not be significant to the relative off-rates that only depend on the pH-switch displayed by CoV1 but not CoV2 (Fig. 3).

**Table 2.** Free energy estimates and baseline rate of detachment of RBD/ACE2 complex. Experimental equilibrium binding free energies are from Walls, *et al*13. Computational free energies 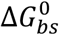 for bound-state (bs) and transition-state (ts), which discounts contributions from amino acids in the switching loop 460-475, were calculated using the *FastContact* server and co-crystals shown in Fig. S2 (neutral pH).

| Virus | $\Delta G_{bs}^{exp}$<br>[kcal/mol] | $\Delta G_{bs}^0$<br>[kcal/mol] | $\Delta G_{ts}^0$<br>[kcal/mol] | $\Delta \Delta G_{slip}^0$<br>[kcal/mol] | $K_{off}^0$<br>[s<sup>-1</sup>] | RBD/ACE2<br>bond |
| --- | --- | --- | --- | --- | --- | --- |
| CoV1 | -11.1 | -11.3 | -8.9 | -2.6 | 1.2×10<sup>4</sup> | slip |
| CoV1 | N/A | -8.9 (*ts*) | 0 | -8.9 | 0.3 | - |
| CoV2 | -11.7 | -11.7 | 0 | -11.7 | 0.003 | tight |

## Optimal dwelling times and endocytosis

Based on typical dimensions of cell and virus radius of 12 and 0.12 *μm*, respectively, ACE2 concentrations in the range of 1000-to-10,000 receptors per cell yield an average separation between receptors *D*~0.6 − 0.2 *μm*. Lateral protein diffusion in cell membranes is length-scale dependent, varying between^23^ 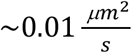 and^24^ 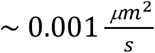 for 40-100 *nm* and >100 *nm*, respectively. Thus, diffusion time scales to bring two receptors into close proximity for the above length scales are ~200 − 5 *s*. On the other hand, the diffusion of the virus in the periciliary fluid can be estimated based on the Stokes-Einstein equation to be 1000 times greater, i.e., 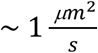, diffusing 1 *μm* in 0.5 s.

Tensile forces triggered by the shear stress in the respiratory airways shorten the lifetime of the CoV1 RBD/ACE2 bond at physiological pH to about 3 s (Table 2), i.e., CoV1 particles bound to cell surface receptors should detach in time scales that are much faster than the time needed for other receptors to bind and stabilize virus attachment. This observation implies that CoV1 will not be able to efficiently engage ACE2 receptors in the upper respiratory tract since after each detachment particles will drift due to normal breathing and gravity down the respiratory tract or be exhaled out (see Supplementary Movie 2). On the other hand, deletion of the pH switch allows CoV2 to have a tighter RBD/ACE2 bond with dwelling times of about ~ 300 s (Table 2), commensurate with the diffusion time scales needed to recruit other ACE2 receptors to stabilize virus attachment (see Supplementary Movie 3).

## Tight-binding at neutral pH found in pangolin-CoV

Further supporting the observation that CoV2 is unique among other coronaviruses, Table 1 compares sequence alignments of pH domains in RBDs of both CoV1, CoV2, MERS, as well as other closely related zoonotic viruses. CoV2-related coronaviruses in pangolin and bats specific strains are missing the pH-switch present in CoV1; instead, they share the Lys458 stabilization motif. However, these zoonotic viruses still have pH-switches that co-localize with the pH-switch in the RBD of MERS (Fig. 1).

Interestingly, as for CoV1, pangolin-CoV species are also one mutation away from losing the last Histidine residue in their corresponding RBD. In order to investigate whether this histidine plays a similar role as His445 in CoV1, we replicated the analysis performed for the RBD of CoV1 in simulations of a homology model of the RBD of pangolin-CoV, and compared the MDs to the co-crystal structure of CoV2 that given their high sequence similarity we used as template for our model (see Methods). Strikingly, we found that neutral (Fig. 4A) and protonated His498 (Fig. 4B) in pangolin-CoV resulted in almost identical phenotypes as for CoV1 but in reverse. Namely, the dynamics of the RBD at neutral pH for NδH consistently displays a bound-like phenotype over the whole simulation time, the same was observed for NεH (Fig. S3). On the other hand, protonation of His498^+^ activates a not-bound-like dynamics in the same switching loop discovered in CoV1 (Fig. 2A), with bound-like behavior for the remaining binding interface (Fig. 4B).

**Figure 4.**
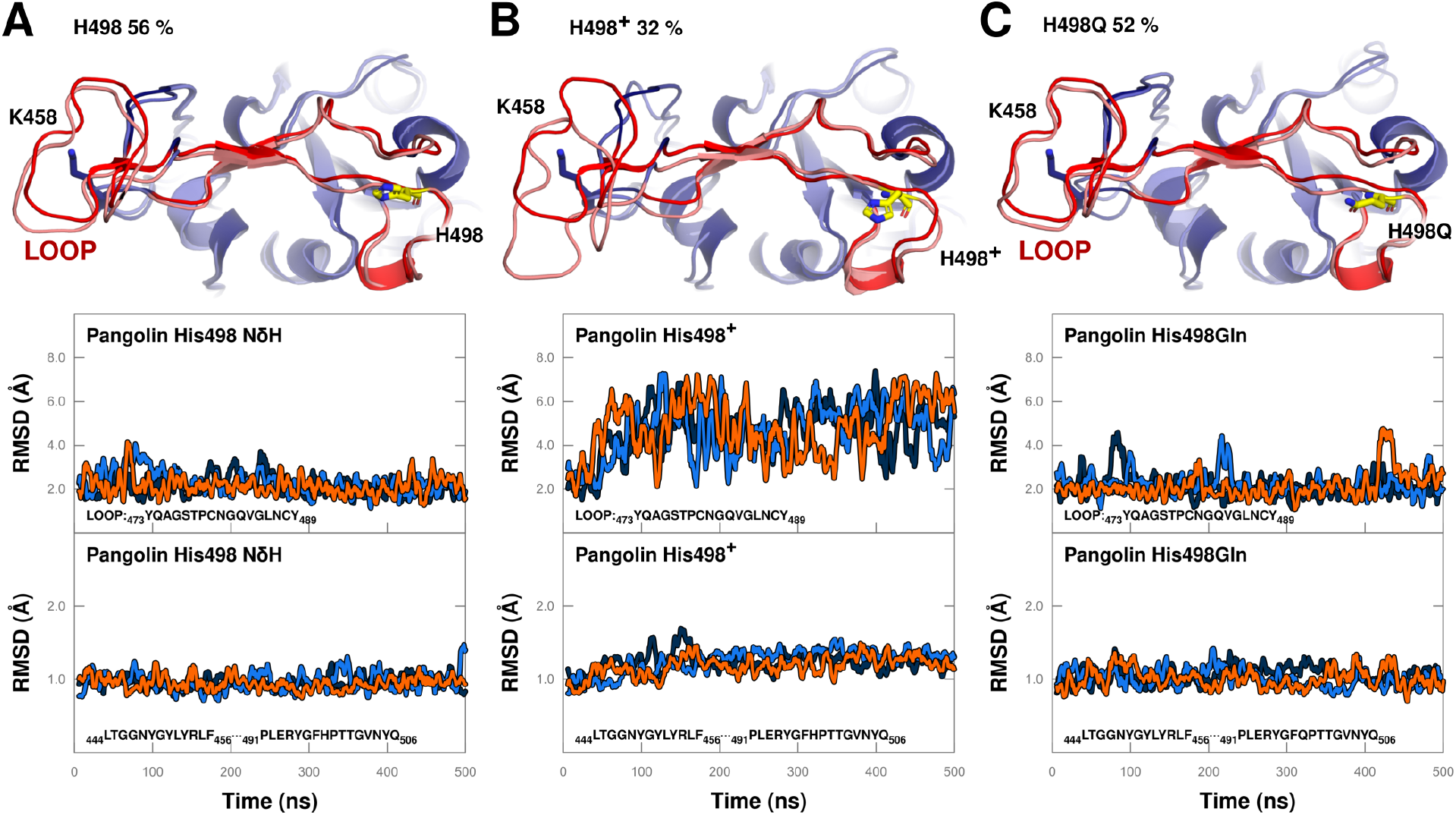
Role of pH switch in binding interface of pangolin-CoV and H498Q mutant. Caption is equivalent as in Fig. 2. Dynamics of three independent MDs show that, at neutral pH, (**A**) pangolin-CoV stays in a bound-like conformations for 100% of the simulation time, and the single point mutation (**C**) His498Gln preserves the same bound-like binding phenotype. (**B**) RMSD of 16 amino acid loop shifts to not-bound-like (~ 6 Å) at low pH (His498^+^).

The observation that, at neutral pH, pangolin-CoV has the same phenotype as CoV2 led us to test a single point mutation at His498 in pangolin-CoV that corresponds to Gln498 in the CoV2 sequence (Table 1). Significantly, we found that this mutation, which makes the RBD of pangolin-CoV pH-independent, is enough to retain the same tight-binding phenotype that we found applies for CoV2, with the dynamics of the whole interface becoming consistently bound-like (Fig. 4C).

## Viruses adopt different mechanisms for cell entry

Molecular dynamics of unbound and bound complexes of the RBD of CoV1 and its specific receptor ACE2 revealed that a pH-switch in the RBD of CoV1 stabilizes a transition state that enhances the off-rate of the RBD/ACE2 bond. In conjunction with the higher diffusion of viral particles relative to cell surface receptors, this “stick-and-slip” mechanism allows the virus to efficiently search the cell surface for random clusters of receptors to stabilize virus attachment and trigger endocytosis in the lower respiratory tract. In principle, “bouncing around” trying to stick to a suitable receptor is also a powerful approach for respiratory viruses to evolve bonds with receptors that are found in abundance in epithelial cell surfaces such as sialic acids for influenza and the ACE2 receptor for SARS (See Supplementary Movie 2).

The newly discovered deletion of the pH-switch in the RBD of CoV2 and its impact on receptor binding reveals a different mechanism (See Supplementary Movie 3) that allows CoV2 internalization to take advantage of the high expression of ACE2 in the nasal epithelium by holding to ACE2 receptors for about 100 times longer than CoV1.

## Viral infection and its pathology

The stick-and-slip molecular mechanism is consistent with CoV1 being mainly a lower respiratory tract disease^25,26^. And, the tighter-bond evolved by CoV2 agrees with recent experiments^1^ that showed robust replication of CoV2 in the primary nasal cells compared to peripheral lung.

Viral replication in human mucous gland cells should release viruses back into the same area^27^ where they can infect new cells until the supply of ACE2 receptors is depleted below the critical threshold needed for binding and internalization. This process will trap viral particles in the upper respiratory tract, naturally leading to longer incubation times. Similarly, accumulation of viral particles in the nasal mucosa will lead to extended periods of viral shedding.

Based on our findings, the distinguishable feature of CoV2 having a higher retention rate in the nasal cavity of adults is a higher transmission rate than CoV1. Interestingly, children do not have well developed sinuses until adolescence^28^. Thus, large areas for viral retention and replication will not be available in young children, which might result in shorter incubation times due to the faster diffusion down to the lower respiratory tract. Something similar could apply to females who have smaller nasal cavities relative to males^29^. Shorter times in the nasal cavity would lead to a lower viral load in the upper airways and could explain the lower transmission observed in young children. Moreover, in light of the new findings by Hou et al.^1^, a weaker replication in this area might impact disease prognosis.

## Evolutionary origin of SARS-CoV2

While we have not yet found the species or strain where the loss of the pH-switch first occurred, relationships shown in Table 1 point at the possible zoonotic origin of CoV2. In particular, the recent finding that the RBD of coronaviruses in tropical pangolins seized in China three years ago^1122^ share a high sequence similarity with the RBD of CoV2 suggests pangolins as a host.

Mechanistically, we found that different pH-switches in SARS-CoV-like viruses regulate the dynamics of the same loop between bound-like and not-bound-like conformations. While pangolin-CoV has the same Lys458 motif of CoV2 that eliminated the pH-switch present in CoV1 at His445, a different pH-switch at His498 stabilizes the bound-state of the RBD at neutral pH, but not at low pH as the CoV1 pH-switch did. However, a single point mutation His498Gln in pangolin-CoV not only eliminates the pH-switch but also retains the bound-like conformations of the RBD operational at both neutral and low pH, i.e., the same tight binding phenotype as that observed in CoV2. This molecular mechanism reveals a direct functional link that supports coronaviruses in pangolins species as the zoonotic origin of CoV2.

## Outlook

While the evolutionary advantage of higher infectivity by SARS-CoV2 in the nasal area is clear, this property comes at the expense of an important regulatory mechanism. Namely, simply based on diffusion, the life-cycle of SARS-CoV2 is significantly slower than that of SARS-CoV because CoV2 is essentially immobilized at its initial cell receptor contact.

The pH-switch in SARS-CoV, which originated in temperate Asia, stabilizes a transition state that enhances the off-rate from its receptor. On the other hand, Sunda pangolins, which originate in a tropical climate, show a tighter bond to ACE2 at neutral pH. These observations suggest that coronaviruses of tropical pangolins smuggled to temperate Wuhan could have undergone adaptive relief in adopting a different binding strategy to their receptors, highlighting the need to better understand the consequences of niche shifts driven by global movement of virus strains.

Collectively, our studies provide insight pertinent to the molecular basis of viral infectivity and, at the same time, validate this form of thermodynamic and molecular modeling as an approach to probe the evolution of the next SARS-mediated pandemic. From a therapeutic perspective, our findings linking viral pathology with long-term viral infection/retention in nasal epithelium of the upper respiratory tract suggest that vaccine development should not just concentrate on fighting systemic infection through induction of IgG responses, but should instead aim to elicit high titers of secretory IgA antibodies capable of neutralizing the virus in the nasal mucosa. Therefore, intranasal delivery of a vaccine with strong IgA producing potential is a logical approach to consider as the next step in countering the current and future pandemics that, like SARS-CoV2, export evolutionarily successful strains via higher transmission rates.

## Methods

Atomic coordinates for starting structures were acquired from the Protein Data Bank^1^: 2DD8 was used for CoV1 RBD and 6LZG was used for the CoV2 RBD. The RBD from 2DD8 (bound to neutralizing antibody) was chosen instead of that in PDB ID 2AJF (bound to ACE2) as a starting structure as it includes otherwise missing portions of the domain. Modification of His to different tautomeric or protonation states was done with PyMol’s Mutagenesis Wizard.^2^ Molecular dynamics simulations (MDs) were carried out with pmemd.cuda from AMBER18^3–5^ using AMBER ff14SB force field^6^ and Generalized Amber Force Field (GAFF).^7^ We used tLeap binary (part of AMBER18) for solvating the structures in a cubed TIP3P water box with a 10 Å distance from structure surface to the box edges, and closeness parameter of 0.75 Å. The system was neutralized and solvated. Simulations were carried out after minimizing the system, gradually heating the system from 0 K to 3 00K over 50 ps, and equilibrating the system for 1 ns at NPT. 500 ns of production was then carried out using NPT at 300 K with the Langevin thermostat, a non-bonded interaction cut off of 8 Å, time step of 2 fs, and the SHAKE algorithm to constrain all bonds involving hydrogens.

Clustering was completed using CPPTRAJ^8^ and H-bond and RMSD calculations were done with VMD.^9^

Homology models of RBD of pangolin-CoV were done making point mutations using PyMOL in CoV2 RBD from PDB 6LZG.

All figures were drawn using PyMOLand GNUPLOT.

## Acknowledgements

This work was supported by NIH GM097082, NS043277 and T32EB009403. CJC is grateful to Dr. Jose Padial for his inspiration in starting this project, to Dr. Dana Ascherman for helping in the immunobiology component of the model, to Drs. Marc Herant and Harinder Singh for successfully challenging earlier interpretations of the model, to Dr. AR Carvunis for highlighting evolution, and to Dr. Olja Finn for her enthusiasm for our model and helpful suggestions.

## Author Contributions

CJC developed the mechanistic ideas and structural modeling, and wrote the paper. KAP and US performed and analyzed the molecular dynamics simulations.

## Additional Information

**Supplementary information** is available for this paper

**Correspondence and requests for materials** should be addressed to CJC at

## Supplementary Information

**Figure S1.**
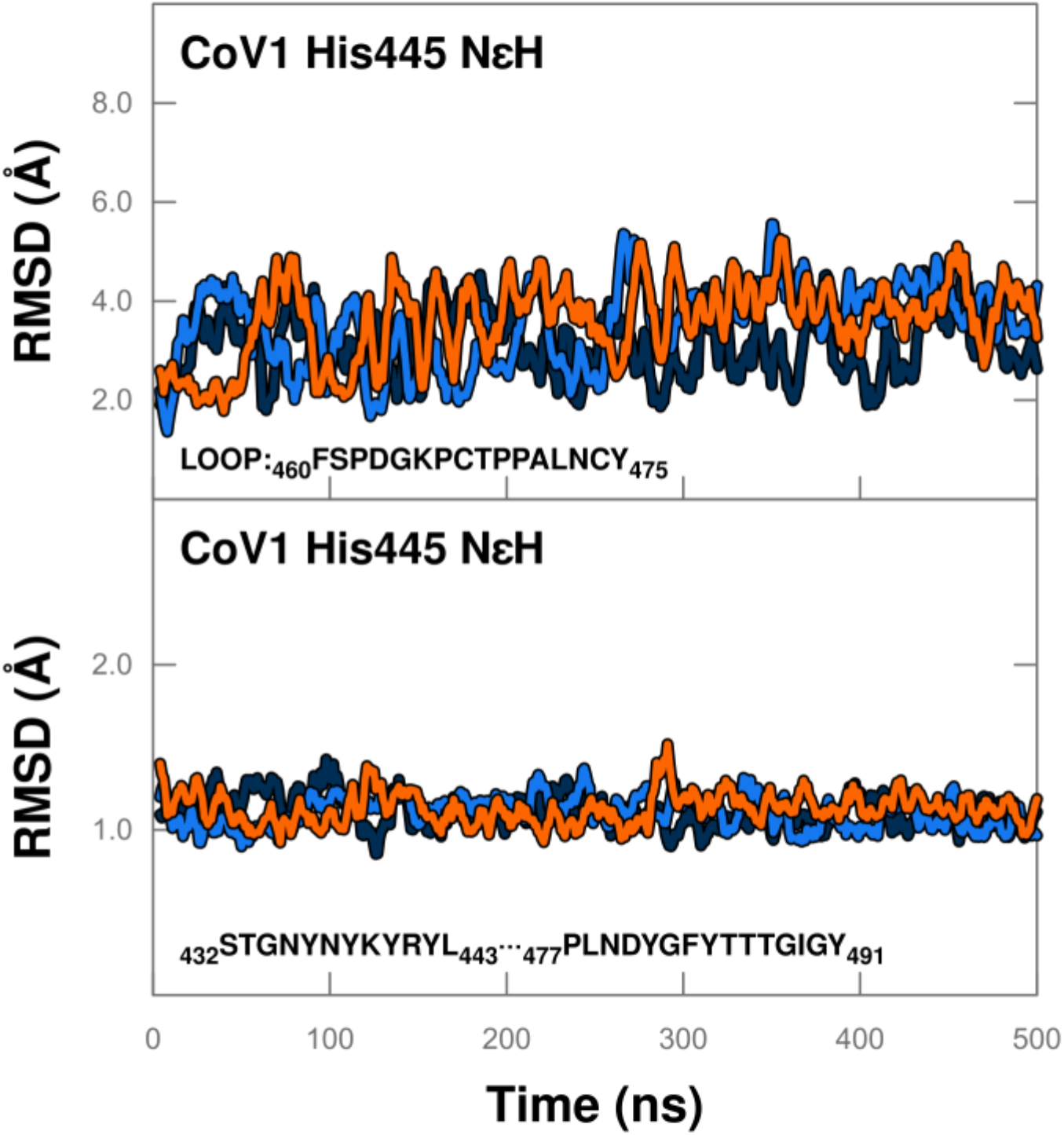
Three independent molecular dynamics of RBD of CoV1 with tautomer NεH at His445. RMSD relative to co-crystal of (a) 16 amino acid loop as a function of time samples not-bound-like states (~4 Å), and (b) the remainder of the binding interface that stays in a bound-like conformation for 100% of the simulation time,

**Figure S2.**
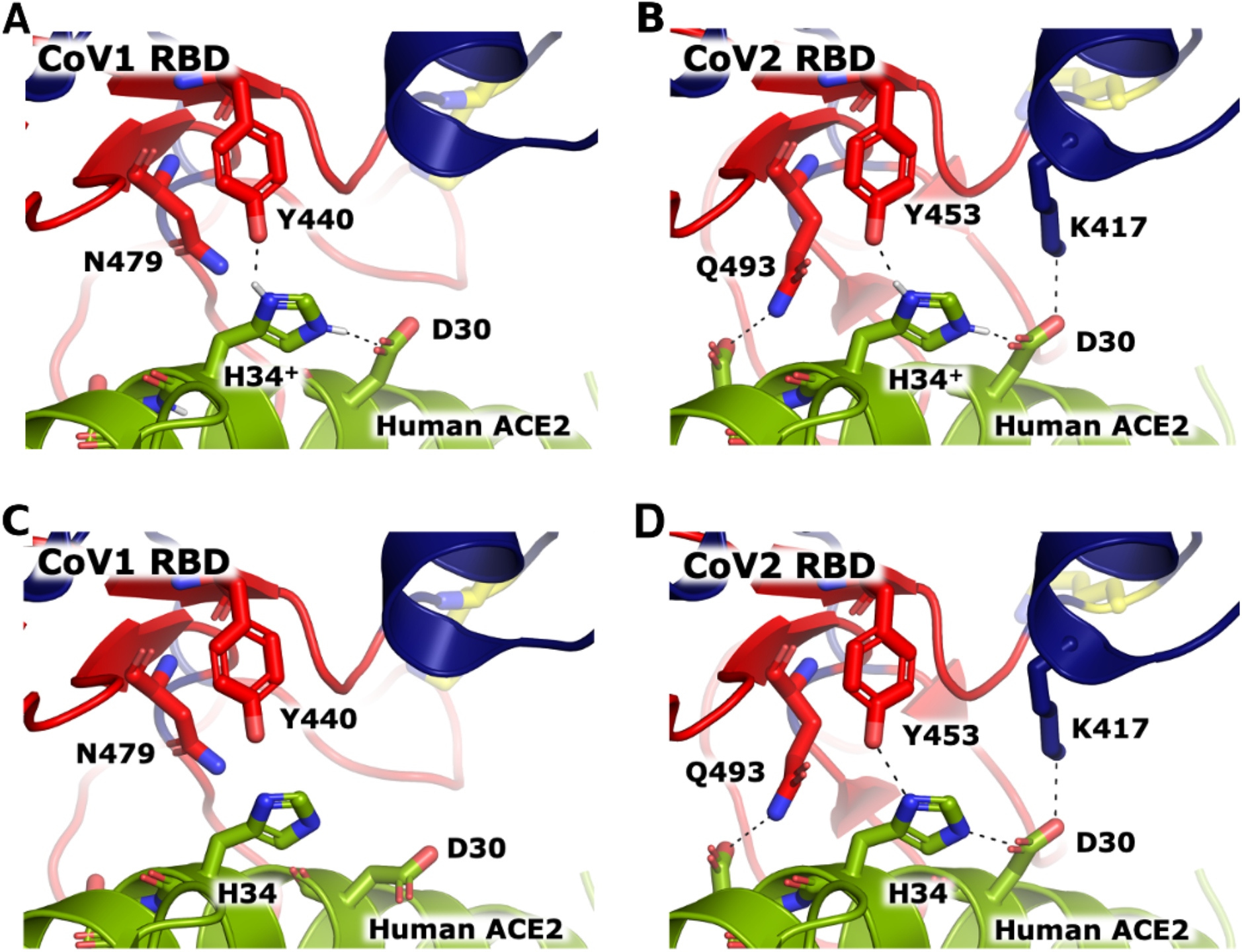
Protonation of the Human ACE2 H34 further stabilizes binding interface. Protonated state of H34^+^ is predicted to form the same stable H-bond network with D30 in ACE2, and Y440 and Y453 in (A) CoV1 and (B) CoV2, respectively. H-bond network is based on rotamers already observed in co-crystals of the unprotonated forms: (C) PDB 2AJF for CoV1, and, (D) PDB 6LZG for CoV2 (see also PDB 6M0J). Unprotonated co-crystal structures of CoV2 assigned the tautomer of H34 to be NδH making a bond with backbone oxygen of D30, which is already making a bond in the α-helix. Our MDs suggest that even in the unprotonated form the rotamer should be rotated 180° having Nδ and Nε interacting with D30 and Y453 (as shown in panels C and D).

**Figure S3.**
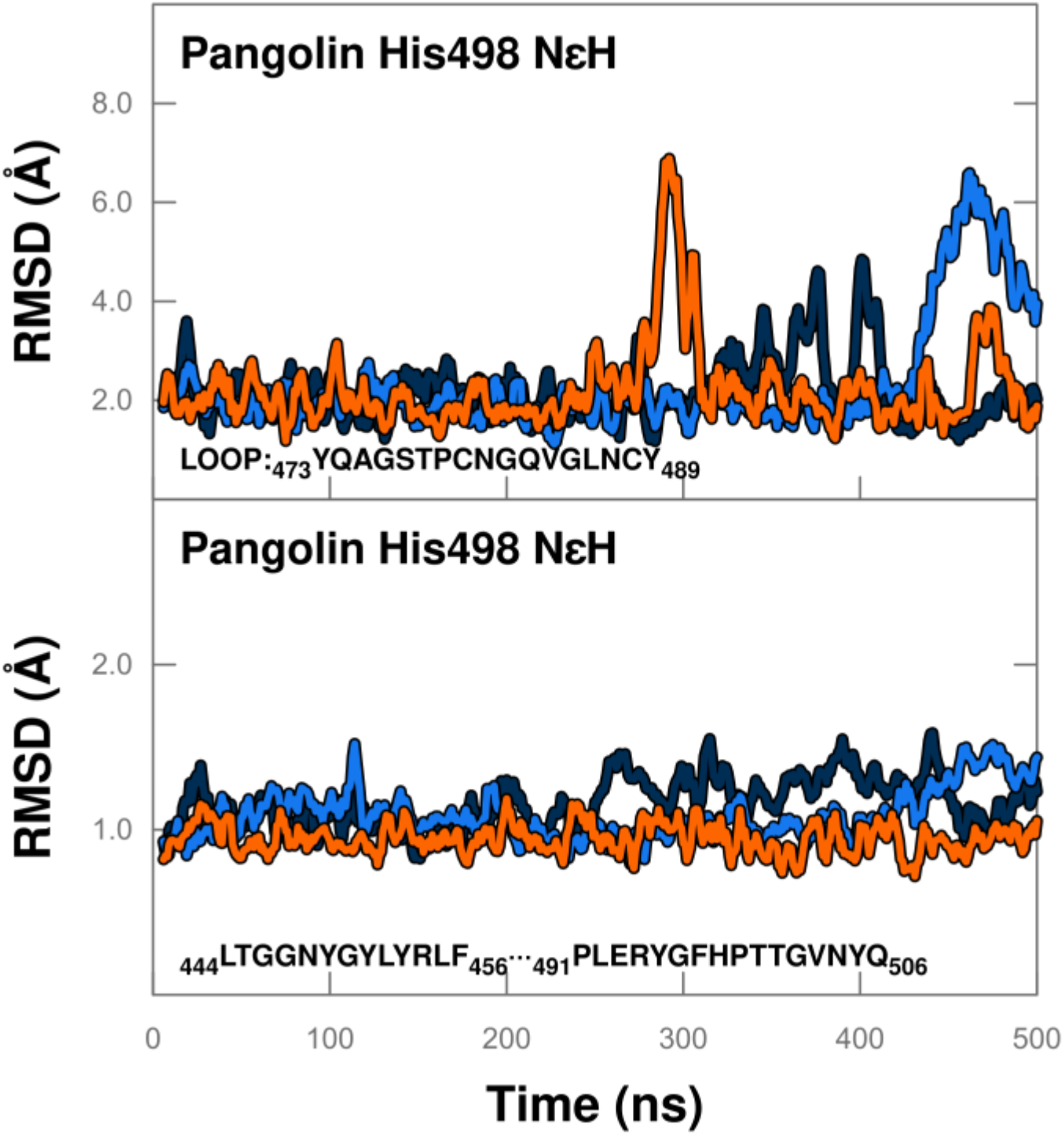
Three independent molecular dynamics show bound-like behavior of Pangolin-CoV with tautomer NεH at His498. RMSD relative to co-crystal of (a) 16 amino acid (homologous) loop, and (b) the remainder of the binding interface as a function of time.

## Supplementary Video 1A

**Title: Representative unconstrained molecular dynamics simulation of CoV1 RBD (PDB 2DD8) at physiological pH (neutral) reveals sampling of a not-bound-like ensemble of the binding interface loop.**

Bound crystal structure 2AJF is shown in dark blue, MD is shown in cyan with the binding interface, including the loop (F_460_SPDGKPCTPPALNCY_475_), in red. ACE2 is depicted in green. H445 in the RBD is in yellow sticks while H34 of ACE2 is shown in dark blue sticks.

## Supplementary Video 1B

**Title: Representative unconstrained molecular dynamics simulation of CoV1 RBD (PDB 2DD8) at low pH reveals sampling of a bound-like ensemble of the binding interface loop.**

Bound crystal structure 2AJF is shown in dark blue, MD is shown in cyan with the binding interface, including the loop (F_460_SPDGKPCTPPALNCY_475_), in red. ACE2 is depicted in green. H445^+^ in the RBD is in yellow sticks while H34 of ACE2 is shown in dark blue sticks.

## Supplementary Video 1C

**Title**: **Representative unconstrained molecular dynamics simulation of CoV2 RBD (PDB 6LZG) reveals sampling of a bound-like ensemble of the binding interface loop.**

Bound crystal structure 6LZG is shown in dark blue, MD is shown in cyan with the binding interface, including the loop (Y_473_QAGSTPCNGVEGFNCY_489_), in red. ACE2 is depicted in green. K458 in the RBD is in yellow sticks while H34 of ACE2 is shown in dark blue sticks.

## Supplementary Video 2

**Title: SARS-CoV-1 virus particle sticks-and-slips from cell surface receptors in upper respiratory tract**

## Supplementary Video 3

**Title: SARS-CoV-2 virus particle binds tightly to cell surface receptors in upper respiratory tract**

